# Dynamic analysis of sequestration-based feedbacks in cellular and biomolecular circuits

**DOI:** 10.1101/2022.03.26.485894

**Authors:** Supravat Dey, Cesar A. Vargas-Garcia, Abhyudai Singh

## Abstract

Nonlinear feedback controllers are ubiquitous features of biological systems at different scales. A key motif arising in these systems is a sequestration-based feedback. As a physiological example of this type of feedback architecture, platelets (specialized cells involved in blood clotting) differentiate from stem cells, and this process is activated by a protein called Thrombopoietin (TPO). Platelets actively sequester and degrade TPO, creating negative feedback whereby any depletion of platelets increases the levels of freely available TPO that upregulates platelet production. We show similar examples of sequestration-based feedback in intracellular biomolecular circuits involved in heat-shock response and microRNA regulation. Our systematic analysis of this feedback motif reveals that platelets induced degradation of TPO is critical in enhancing system robustness to external disturbances. In contrast, reversible sequestration of TPO without degradation results in poor robustness to disturbances. We develop exact analytical results quantifying the limits to which the sensitivity to disturbances can be attenuated by sequestration-based feedback. Next, we consider the stochastic formulation of the circuit that takes into account low-copy number fluctuations in feedback components. Interestingly, our results show that the extent of random fluctuations are enhanced with increasing feedback strength, but can exhibit local maxima and minima across parameter regimes. In summary, our systematic analysis highlights design principles for enhancing the robustness of sequestration-based feedback mechanisms to external disturbances and inherent noise in molecular counts.

## I. Introduction

As in engineering, feedback regulation forms the key basis of homeostasis in physiological and cellular systems. Perhaps the simplest example of this is seen in the regulation of gene expression, where a transcription factor binds to the promoter of its own gene to modulate transcriptional activity. Such forms of gene autoregulation are key motifs in gene regulatory networks [1], and have been thoroughly investigated using a combination of mathematical and experimental tools. These studies highlight how gene autoregulation can alter response times [2]–[4], attenuate the impacts of stochasticity [5]–[18], and alter the system’s information capacity [19]. There is a growing appreciation of similar autoregulatory feedbacks at play in cell populations controlling their density through extracellularly secreted factors [20]–[23], and neurotransmitter binding to autoreceptors on neuronal membranes to maintain homeostatic activity [24]–[27].

In this study, we go beyond autoregulation to consider a nonlinear feedback system that we refer to as *sequestration-based feedback.* It can represent by the following system of chemical reactions

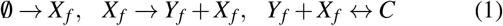

where a species *X_f_* is produced constitutively at a constant rate and activates the production of species *Y_f_*. *Y_f_* reversible binds to *X_f_* sequestering it in a complex *C*. In essence, an increase in *Y_f_* levels results in a downregulation of its synthesis through sequestration of its activator *X_f_*. Several examples of this can be seen in biology as illustrated in Fig. 1. As a physiological example, hematopoietic stem cells residing in the bone marrow are induced to differentiate into platelets (*Y_f_*) by a protein called Thrombopoietin (*X_f_*). However, megakaryocytes (precursor cells that produce platelets) and platelets actively sequester Thrombopoietin (TPO) and degrade it. Thus, depletion of platelets in response to injury leads to an upregulation in platelet synthesis via an increase in free TPO levels [28]–[30]. While the differentiation of stem cells into platelets is a complex process occurring through several intermediate states, for modeling simplicity we consider platelet synthesis occurring in a single TPO-dependent rate. A key focus is to understand how platelet-induced degradation of sequestered TPO can enhance the system’s robustness to fluctuations in platelet demand.

**Fig. 1.**
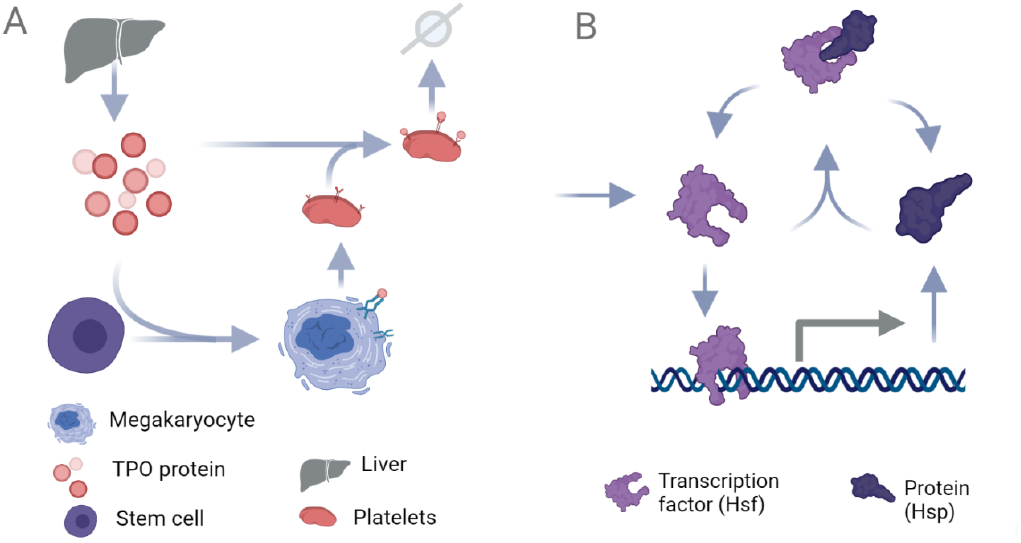
Examples of sequestration-based feedback in physiology and cellular responses. A: The differentiation of stem cells into megakaryocytes and platelets is driven by a protein called Thrombopoietin (TPO) that is produced from the liver. Megakaryocytes/platelets sequester and degrade TPO implementing a feedback mechanism where depletion of platelets increases their synthesis via an upregulation in freely available TPO [28]–[30]. B: The biomolecular circuit orchestrating the cellular heat-shock response. The heat-shock protein (Hsp) binds and sequesters the heat-shock factor (Hsf), and free Hsf proteins upregulate the synthesis of Hsp.

A classic biomolecular example of sequestration-based feedback is the cellular heat-shock response (Fig. 1B), where heat-shock protein (*Y_f_*) sequesters the heat-shock factor (*X_f_*). Exposure to heat denatures proteins, and heat-shock proteins (Hsp) are chaperones that assist in protein refolding. The binding of Hsp to client proteins results in an increase in free heat-shock factor (Hsf) levels as there is less free Hsp available to bind to Hsf. Hsf is a transcription factor (TF) that once released from the complex binds to the Hsp gene promoter to upregulate its synthesis closing the feedback loop [31]–[33].

Other examples of sequestration-based feedback include biomolecular circuits involved in controlling rDNA repeats [34] and the growing evidence of feedback between specific microRNAs and TFs. In the latter example, TFs activate the synthesis of microRNAs and the feedback is implemented by having the microRNA bind and sequester the TF mRNA to turn off TF synthesis [35], [36]. It is important to point out that a special case of sequestration-based feedback is the antithetic integral feedback that has been recently studied [37]–[40] and also experimentally implemented via synthetic circuits [41], [42]. Such forms of integral feedback play a vital role in the adaptation of biological processes to changes in input stimuli [43]–[46].

Overall, the paper is organized as follows. In Section II, we develop a nonlinear dynamical model for the sequestration-based feedback. We study in Section III how sensitive the total platelet abundances are to fluctuation in platelet consumption rate. Our analysis derives fundamental limits to which this sensitivity can be reduced and finds platelet-induced TPO decay to be critical for robust buffering. In Section IV, we turn to a stochastic formulation of the heatshock response circuit and investigate feedback performance in the presence of inherent stochasticity in gene expression.

## II. Deterministic model formulation

The system of chemical events that implement the sequestration-based feedback is shown in Table I along with the net rates with which these reactions occur per unit time. Here capital letters denotes species:

- *X_f_* – free Thrombopoietin (TPO) not bound to platelets
- *Y_f_* – free platelets not bound to TPO
- *C* – platelets bound to TPO.

**TABLE I.** Different chemical reactions constituting the sequestration-based feedback along with their net rates

| Reaction | Net rate | Comment |
| --- | --- | --- |
| $\emptyset \rightarrow X_f$ | $k_x$ | Synthesis of Thrombopoietin (TPO) |
| $X_f \rightarrow Y_f + X_f$ | $k_y x_f$ | Production of platelets by free TPO |
| $Y_f + X_f \rightarrow C$ | $k_b x_f y_f$ | Binding of TPO to platelets |
| $C \rightarrow Y_f + X_f$ | $k_u c$ | Unbinding of TPO from platelets |
| $C \rightarrow Y_f$ | $\gamma_i c$ | Platelet-induced decay of TPO |
| $C \rightarrow \emptyset$ | $\gamma_y c$ | Degradation of TPO-bound platelets |
| $Y_f \rightarrow \emptyset$ | $\gamma_y y_f$ | Degradation of TPO-unbound platelets |
| $X_f \rightarrow \emptyset$ | $\gamma_x x_f$ | Degradation of free TPO |

The corresponding small letters *x_f_*(*t*), *y_f_*(*t*), *c*(*t*) represent the concentration of species at time *t*. To be physiologically relevant, these state variables can only take positive real values. Combining the rates of production and loss for each chemical species results in the following nonlinear dynamical system describing the time evolution of concentrations

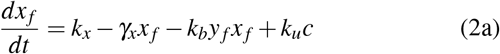

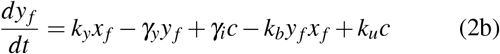

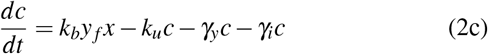

where *k_x_* is the constitutive rate of TPO synthesis from the liver. Free TPO molecules drive the synthesis of new platelets with rate *k_y_x_f_*, and we refer to *k_y_* as the *platelet activation rate.* Binding and unbinding of TPO to platelets occurs with rates *k_b_* and *k_u_*, respectively, as per massaction kinetics. An important parameter representing platelet-induced degradation of bound TPO is *γ_i_* that converts a TPO-bound platelet back into a free platelet (*C* → *Y_f_* reaction in Table I). Later on we will specifically investigate the role of *γ_i_* in platelet homeostasis. Finally, free TPO is degraded with rate *γ_x_*, and *γ_y_* is the rate of consumption of platelets that is assumed to be the same irrespective of whether it’s bound or unbound to TPO. Here 1/*γ_y_* is average lifespan of an individual platelet that has been measured to be ≈ 1 week in humans [47]–[49].

Since we are primarily interested in the total amount of platelets, (2) is transformed to

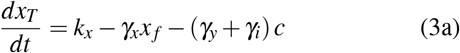

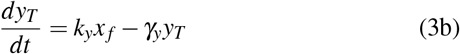

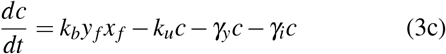

by considering a change of variables

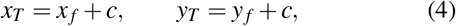

where *x_T_* and *y_T_* denote the total (bound plus unbound) level of TPO and platelets, respectively. In the limit of fast binding/unbinding (i.e., *k_u_* →> ∞ and *k_b_* → ∞) for a given dissociation constant

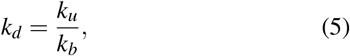

the free platelet and TPO concentrations rapidly equilibrate such that

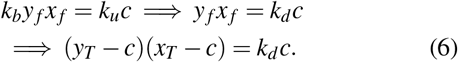

Thus, given the total platelet and TPO levels (*y_T_* and *x_T_*) at any time instant, solving (6) yields the bound platelet level

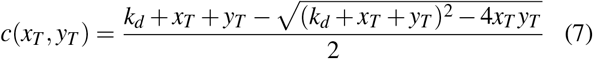

resulting from fast binding and unbinding processes. Before proceeding further, we highlight key properties of this continuously differentiable function *c*(*x_T_, y_T_*), ∀*x_T_, y_T_* ≥ 0:

- *c*(0, *y_T_*) = 0 and *c*(*x_T_*, 0) = 0.
- 0 ≤ *c*(*x_T_, y_T_*) ≤ Minimum of *x_T_* and *y_T_*.
- *c*(*x_T_, y_T_*) monotonically increases with respect to both arguments with bounded slope (proof in Appendix):

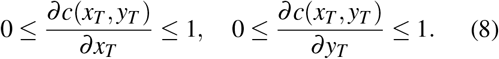
- In the limit of rapid TPO dissociation (i.e., no binding of TPO to platelets)

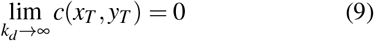

Exploiting this timescale separation, the original model (3) is now reduced to a two-dimensional model

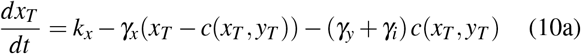

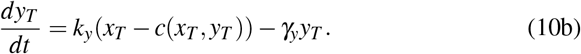

Having formulated the model we present our first result on the uniqueness and stability of its equilibrium point.

### Theorem

The two-dimensional nonlinear system (10) has a unique *asymptotically stable* equilibrium given by

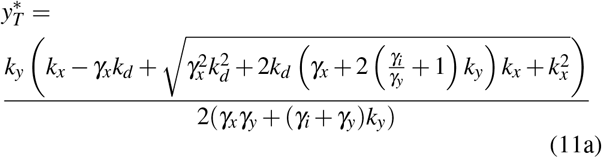

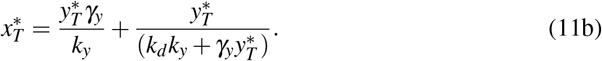

### Proof

Solving the nonlinear system (10) at steady-state yields a unique equilibrium point given by (11). As expected, in the limit of no TPO binding to platelets this equilibrium reduces to

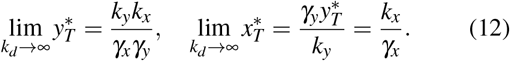

Linearizing the nonlinear systems around (11) yields the following Jacobian matrix

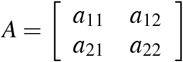

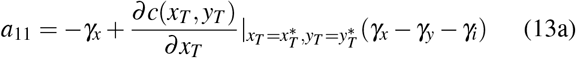

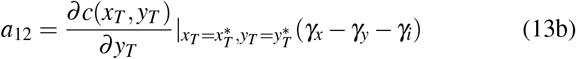

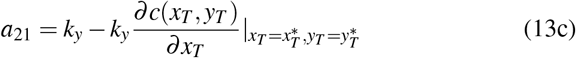

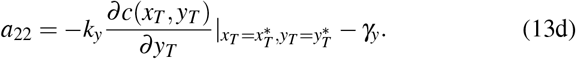

The trace of the *A* matrix is given by

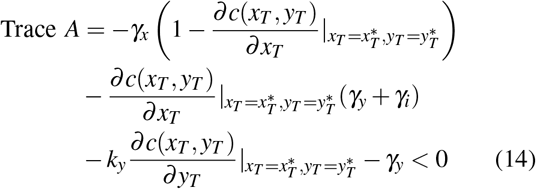

which is strictly negative, as from (8), *c*(*x_T_,y_T_*) is a monotonically increasing function in both arguments with

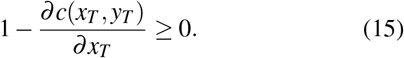

Thus, each of the three terms in (14) are individually negative, making the trace negative. Using a similar argument we can see that all the three terms making up the following determinant of the *A* matrix

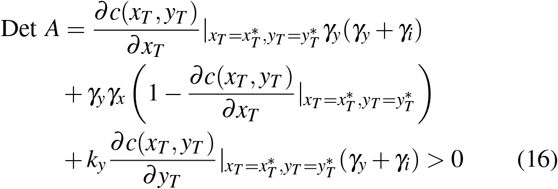

are individually positive, hence implying a positive determinant. For a 2 × 2 *A* matrix, a negative trace and a positive determinant are *necessary and sufficient* for asymptotic stability [50]. We also note that

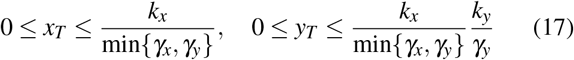

is an invariant set, i.e., trajectories starting outside the set will enter it in finite time and remain there. More specifically, if

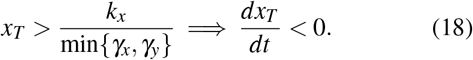

This can be seen by considering two cases, first if *γ_y_* > *γ_x_*, then the right-hand-side of (10a) is

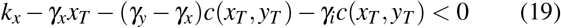

if 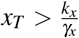. Secondly if *γ_y_* < *γ_x_*, then the right-hand-side of (10a) can be rewritten as

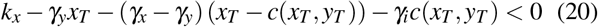

if 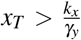 as *c*(*x_T_, y_T_*) ≤ *x_T_*. Once the trajectory reaches 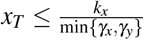 then from (10b) if

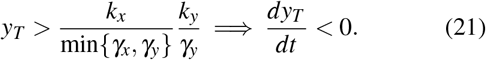

## III. Sensitivity analysis

Having determined the model’s equilibrium, we next study the sensitivity of the total platelet abundance 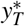 to the consumption rate *γ_y_* which can dramatically go up in response to injury. This can be quantified by the dimensionless log sensitivity

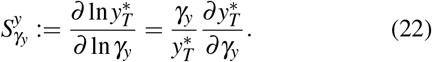

From (11), in the absence of platelet-induce TPO decay (*γ_i_* = 0) the total platelet abundance

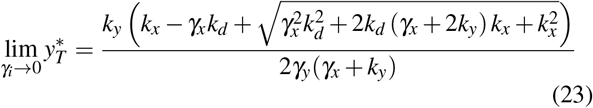

is inversely proportional to *γ_y_* that results in 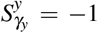. Thus, a hundred-fold increase in *γ_y_* results in a hundred-fold decrease in platelet abundance (Fig. 2). In contrast, presence of platelet-induced TPO decay (*γ_i_* > 0) can significantly buffer this decrease by making 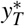 less sensitive to *γ_y_* (Fig. 2). Are their limits to which sensitivity can be reduced and how does this limit depend on *γ_i_*?

**Fig. 2.**
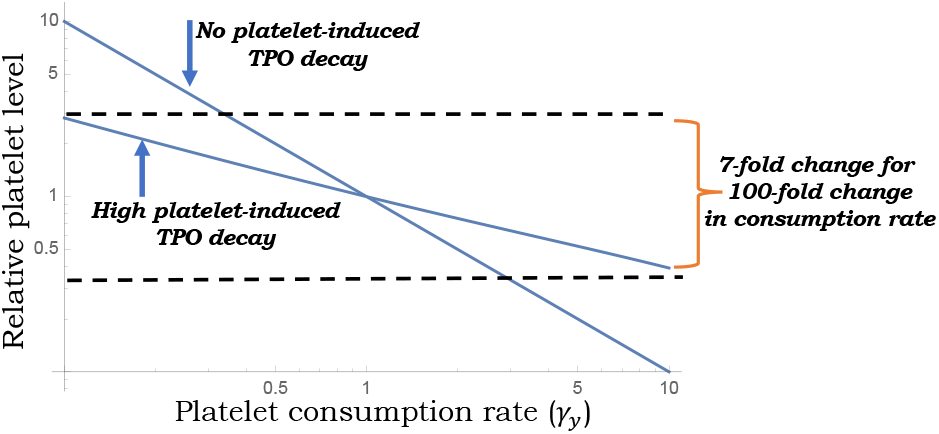
Platelet-induced TPO decay buffers platelet abundance to increase in consumption rates. Plot of total platelet abundance 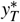 as given by (11) for varying consumption rate *γ_y_*. A hundred-fold change in *γ_y_* results in a hundred-fold (*χ* = 0) and seven-fold (*γ_i_* = 50) change in 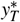. Other parameter taken as *k_y_* = 10, *k_d_* => 1 and *k_x_* was varied so as to have 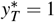 when *γ_y_* = 1. All timescales are normalized to TPO lifespan that results in *γ_x_* = 1, and both TPO/platelet levels have arbitrary units.

To quantify the fundamental limit to which sensitivity can be suppressed, we derive the following general formula for 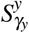 from (11)

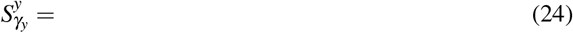

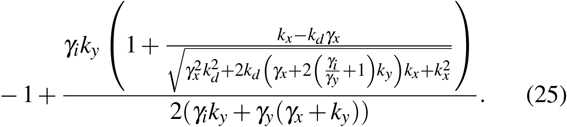

Our analysis of this formula reveals that it can either vary monotonically or non-monotonically with respect to the platelet activation rate *k_y_*. To see this, we first define the dimensionless constant

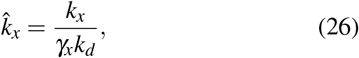

where recall from (12) that *k_x_/γ_x_* is the total TPO level in the absence of any binding to platelets. When 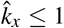, then 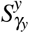 monotonically decreases with increasing activation rate *k_y_* (Fig. 3; top) with the following limits

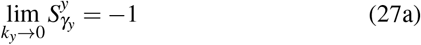

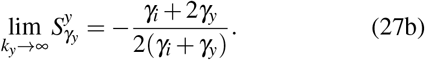

**Fig. 3.**
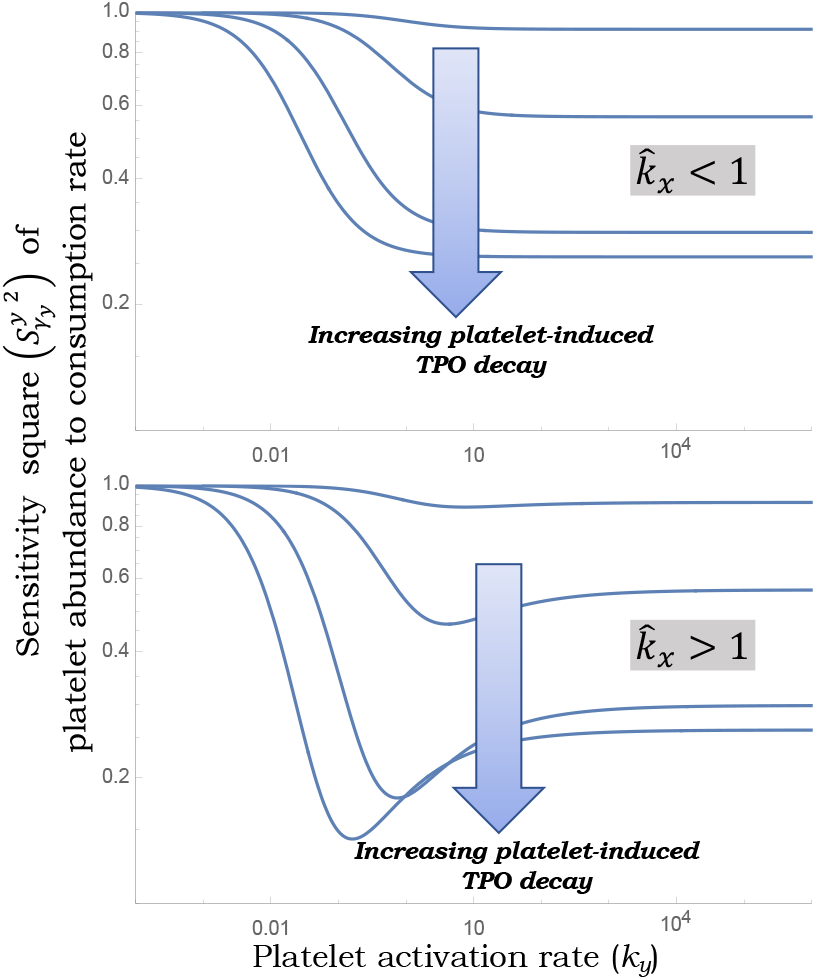
Sensitivity of platelet abundance to consumption rate is minimized at an optimal activation rate. Plot of the square of (24) as a function of *k_y_* for 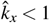 (top) and 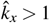 (bottom). Other parameters take an *γ_x_* = *γ_y_* = *k_d_* = 1. The four different curves correspond to *γ_i_* = 0.1,1,10,50.

Thus, in this regime, (27b) gives the fundamental limit of sensitivity repressions with 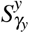 approaching −1/2 as *γ_i_* → ∞.

Interestingly, when 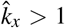, 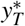 varies non-monotonically with *k_y_* and is minimized at an intermediate value of the platelet activation rate (Fig. 3; bottom). Please note that since 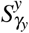 is negative, Fig. 3 plots the square 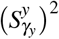. In the case of 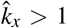, our results show that the optimal reduction of sensitivity occurs when

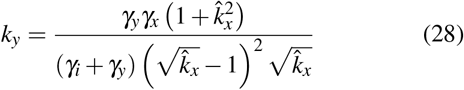

and the corresponding sensitivity is

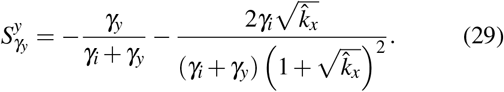

In the limit 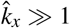, the optimal value of *k_y_* and the corresponding minimal sensitivity simplify to

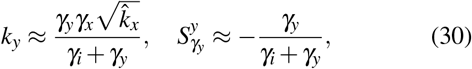

respectively. Note that this limit of sensitivity suppression is inversely related to *γ_i_*, thus can be made arbitrarily small by arbitrarily enhancing the platelet-induced decay rate. The effect of this sensitivity reduction can also be seen in the dynamical response of platelet abundance to a step increase in the consumption rate. (Fig. 4). The optimally chosen value of *k_y_* as per (28) results in a lower depression in *y_T_*(*t*) as compared to higher and lower values of *k_y_* (Fig. 4).

**Fig. 4.**
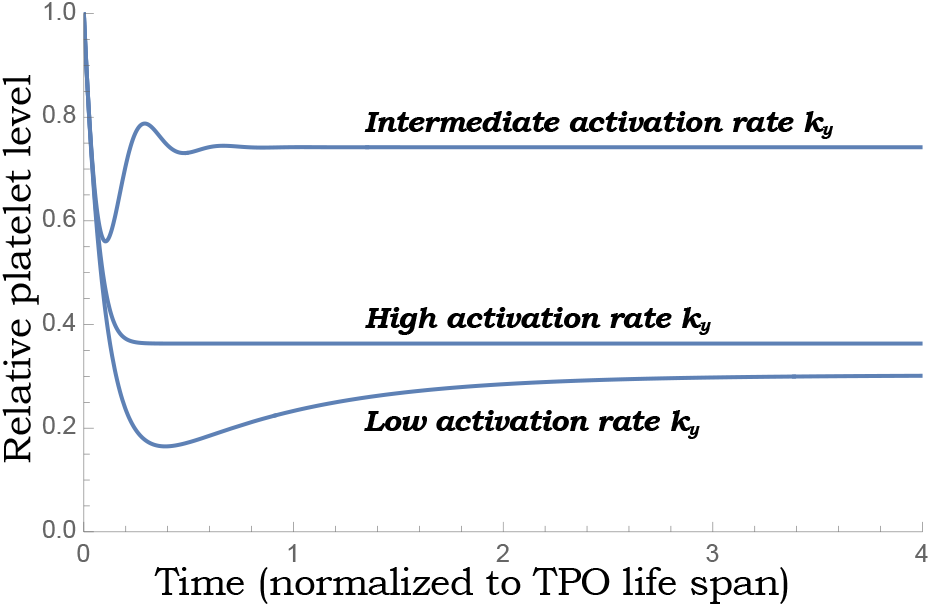
Optimally set activation rate *k_y_* yields minimal depression of platelet count in response to increased consuption. The platelet abundance *y_T_*(*t*) as obtained by solving the nonlinear system (10) for a ten-fold change in *γ_y_* from 1 to 10. Other parameters taken as *χ* = 100, *k_d_* = 0.1, *k_x_* = 100 and *γ_x_* = 1. The intermediate activation rate corresponds to *k_y_* chosen as per (28), and other curves were generated assuming a hundred-fold higher or lower value than (28).

## IV. Biomolecular sequestration-based feedback

We next consider a biomolecular example of sequestration-based feedback as depicted in Fig. 1B. More specifically, the heat-shock protein (Hsp) binds and sequester the heat-shock factor (Hsf), and free Hsf proteins activate the production of Hsp by enhancing its gene’s transcription [31], [32]. The deterministic model formulation is exactly similar to (2) but with redefined species

- *X_f_* – free Hsf protein not bound to Hsp
- *Y_f_* – free Hsp protein not bound to Hsf
- *C* – Hsp protein bound to Hsf.

Moreover, we now refer to *k_y_* and *γ_i_* as the *hsp activation rate* and *hsf-induced hsf decay rate*. Given that these proteins are fairly stable with long half-lives, the decay in their concentrations is primarily from cellular growth during the cell cycle. Towards that end,

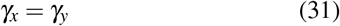

where *γ_y_* is the cellular growth rate. *How robust are total Hsp levels to growth rate fluctuations?*

A formula for sensitivity 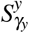 can again be obtained from (11) which is slightly different from that obtained earlier for the TPO-platelet circuit where *γ_x_* and *γ_y_* were independent parameters. However, this formula shows qualitatively similar behavior with

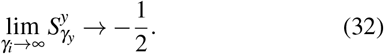

A plot of 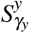 with respect to the activation rate in Fig. 5 shows that as in Fig. 3, for small values of 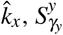 monotonically decreases with increasing *k_y_* to reach the limit (27b). In contrast, for large values of 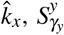 shows a U-shape profile where the minimal sensitivity can be made arbitrarily close to zero by enhancing *γ_i_*. An important quantitate difference to note is that while 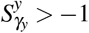 in the TPO-platelet circuit (Fig. 3), in the heat-shock system 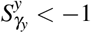 for certain parameter regimes (Fig. 5) implying a decay in 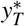 with an increasing growth rate that is even faster than 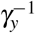.

**Fig. 5.**
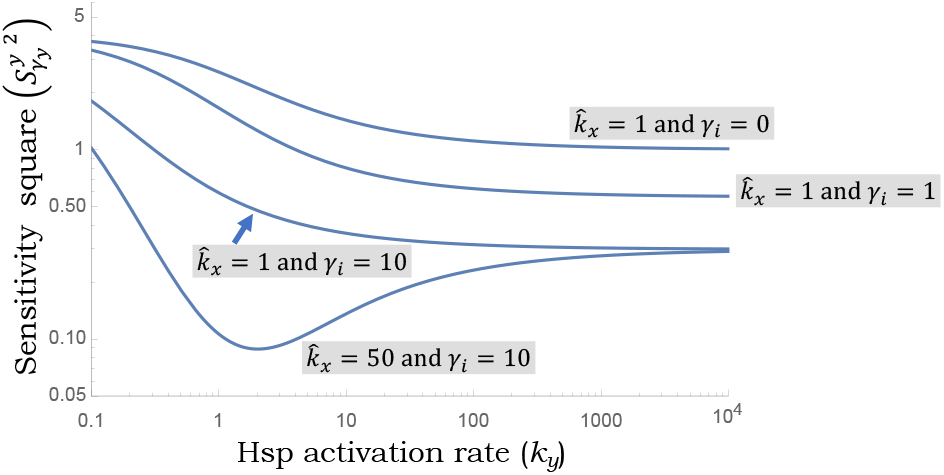
Sensitivity of total heat-shock protein abundance to growth rate is minimized at an optimal hsp activation rate. Plot of the square of the sensitivity 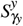 for the biomolecular sequestration-based feedback describing the heat-shock response (Fig. 1B). Here *γ_x_* = *γ_y_*, where *γ_y_* is the cellular growth rate. As in Fig. 3, other parameters are taken as *γ_x_* = *γ_y_* = *k_d_* = 1.

Having examined fluctuations in the growth rate, we next consider an alternative source of stochasticity in the biomolecular circuit that arises from the synthesis of gene products in bursts of transcriptional activity. We analyze the impact of such bursting on total Hsp levels in stochastically-formulated models of sequestration-based feedback.

### A. Stochastic Analysis

We next consider a stochastic formulation of the chemical reaction set in Table I, where Hsf is produced in random bursts. In gene expression, such bursty synthesis is realized due to the short lifespan of mRNA compared to the corresponding protein, as in prokaryotes [51]. In eukaryotes, however, the cause of burst is usually the gene switching between active and inactive states [52]–[56]. Such bursty expressions are the major sources of large fluctuations in expression levels (gene expression noise). As Hsf activates the Hsp production, its noise propagates and makes the Hsp level highly noisy. For simplicity, we assume a simple non-bursty birth process for Hsp production.

#### Notation

In this formalism, *x*(*t*) denotes the number of copies species *X*. It takes a random non-negative integer value i.e., *x*(*t*) ∈ {0,1,2,3,...}. We use angular brackets 〈·〉 and 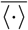 to denote the transient and steady-state expected values for the stochastic process, respectively.

The probability of the Hsf and Hsp production events in infinitesimal interval (*t, t* + *dt*] are given by

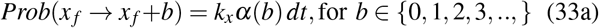

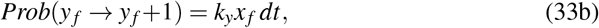

where *α*(*b*) is the burst size distribution, generally follows geometric distributions. The probability of binding and unbinding events are

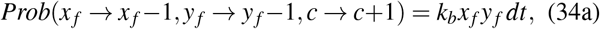

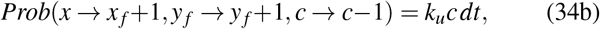

All the species decay with the same rate *γ_x_* = *γ_y_* = *γ_c_*. In addition, there is a Hsp-induced Hsf degradation which occurs at rate *γ_i_*. The probabilities of the degradation events during the infinitesimal interval are

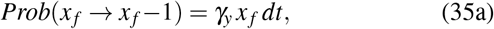

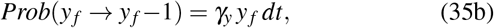

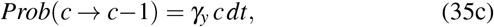

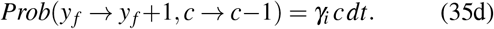

#### Noise in total Hsp count

We are interested in understanding fluctuations in the total Hsp count (*y_T_* = *y_f_* + *c*) at steadystate. The fluctuations, also known as gene expression noise, can be quantified by the Fano factor in *y_T_*

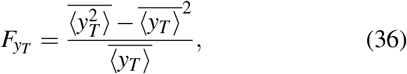

where 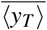 and 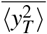 are the first and second-order moments of *y_T_* at steady-state.

The nonlinear binding term makes the exact analytical calculation difficult. In the absence of binding, one can obtain exact analytical results using the chemical master equation framework for discrete-state continuous-time Markov processes [57]. For no binding case,

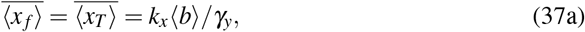

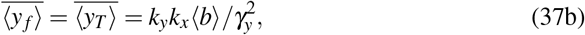

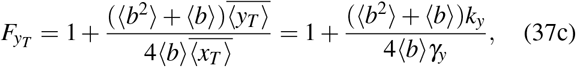

where 〈*b*〉 = ∑_*i*_ *α*(*i*) represents the average burst size and 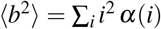. Both 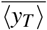 and *F_yT_* linearly depend on the activation rate *k_y_*. The noise level also increases with the average size and width of the burst size distribution.

To analyze the noise in the presence of binding, we perform numerically exact stochastic simulations using the kinetic Monte Carlo method [58]. In Fig. 6, we plot the mean (A) and the noise behavior (B) in the total Hsp count as a function of the activation rate. For very low values of *k_y_* (*k_y_/γ_y_* ≪ 1), 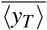 linearly increases with *k_y_*, following the no-binding case above. For an intermediate *k_y_*, the increase in 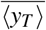 becomes flat, and finally, *k_y_/γ_y_* » 1 the increase becomes faster again but stays well below the no-binding case due to negative feedback (Fig. 6A). The Hsp-induced Hsf decay further reduces 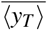 and also decreases the intermediate flat region.

**Fig. 6.**
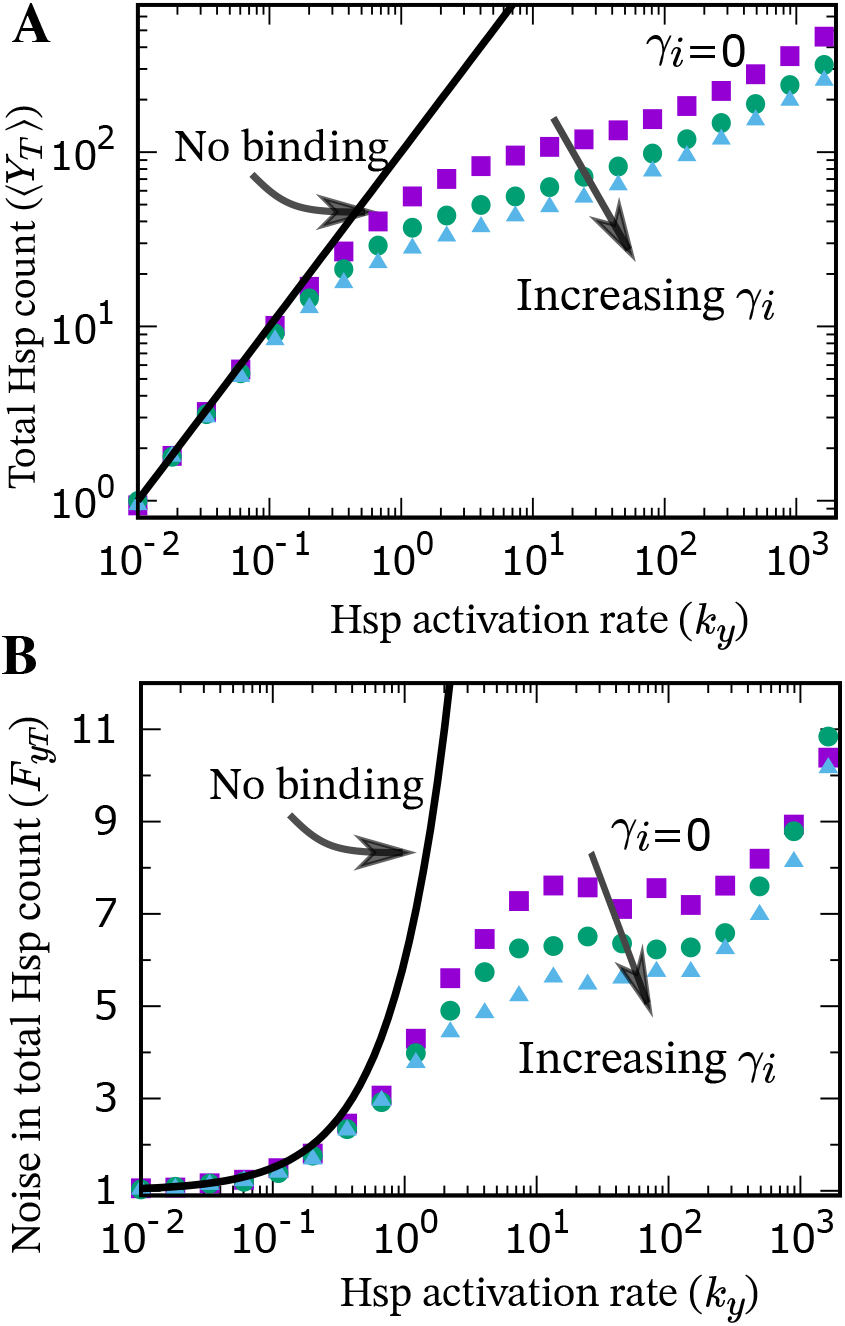
Hsp-induced Hsf degradation reduces noise in total Hsp count. The mean (A) and the Fano factor (B) are plotted against *k_y_* for different values of *γ_i_* (= 0, *γ_y_*, 2*γ_y_*). The lines represent analytical results for no binding case, whereas symbols denote stochastic simulation results in the presence of binding. Parameters used: 〈*b*〉 = 10, *k_x_* = 10, *γ_y_* = 1, *k_d_* = 1 and *k_u_* = 40. Our burst size distribution is a shifted geometric distribution.

Implementing the feedback through the Hsp-Hsf binding reduces the noise in the total Hsp count as compared to the no-binding case (Fig. 6B), and the hsp-induced Hsf decay decreases *F_yT_* further. Interestingly, our results show a nonmonotonicity in *F_yT_* at an intermediate value of *k_y_* when *γ_i_* is relatively small, and this is connected to the flattening in 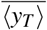 levels seen in Fig. 6A.

## V. Conclusion

There is growing literature on leveraging sequestration processes for feedback control of biomolecular systems [59], [60]. Here we analyzed a sequestration-based feedback architecture illustrated in Table I that mimics homeostatic control occurring at diverse spatial scales from maintaining cell numbers in an organism to regulating intracellular concentrations of key gene products (Fig. 1). Our model presented in (2) reduce to the well-studied antithetic integral feedback in the limit *γ_i_* = *γ_u_* = *γ_y_* = 0 that corresponds to *X_f_* irreversibly binding to *Y_f_*, and both subsequently degrading. In this case (2b) becomes

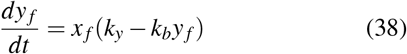

and one can see the integral feedback (albeit with a timevarying coefficient *x_f_*) that regulates *y_f_* around its steadystate value *k_y_/k_b_.* As pointed out earlier [61], from a practical standpoint this integration is “leaky” due to the effects of dilution (*γ_y_* ≠ 0).

Our analysis of the sequestration-based feedback identified limits to which the sensitivity of total *Y_f_* levels to its degradation rate *γ_y_* can be reduced. The key result is presented in (29), where the absolute value of 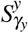 reduces with increasing platelet-induced TPO degradation *γ_i_* reaching the limits

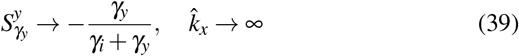

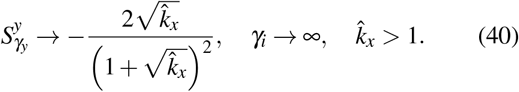

These limits are important for the homeostasis of platelets abundance where there is strong evidence for *γ_i_/γ_y_* » 1 [28]. While there is no evidence for induced decay in the Hsp-Hsf feedback circuit, our analysis shows that in the case of *γ_i_* = 0, sensitivity reduction can occur from −1 to −1/2 (Fig. 5) making 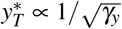.

Analysis of the sequestration-based feedback considering stochasticity of biochemical reactions and transcriptional bursting shows a noise reduction in the feedback system compared to no feedback (i.e., no binding between *X_f_* and *Y_f_*) in the absence of induced decay (*γ_i_* = 0). Importantly, noise further attenuates with increasing *γ_i_* (Fig. 6). While this investigation relies on stochastic simulation of the underlying biochemical reactions, future work will use analytical tools based on moment closure schemes [62]–[64] to systematically investigate noise properties, specifically focusing on the origin of local minima in the Fano factor (Fig. 6).

## Appendix

Taking the derivative of (7) we obtain

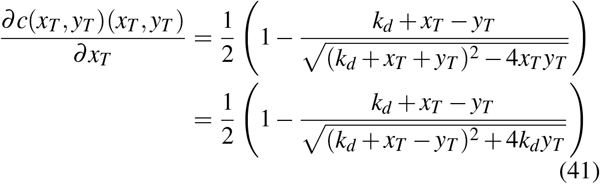

Note that while *k_d_, y_T_, x_T_* only take positive real values, *k_d_* + *x_T_* – *y_T_* can be either negative or positive depending on the total abundances of TPO and platelets. However, irrespective of the sign of *k_d_* + *x_T_* – *y_T_*

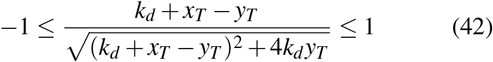

which from (41) implies that

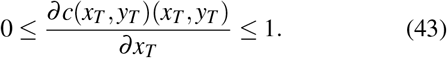

